# Homo-and heterofermentative lactobacilli are differently affected by lignocellulosic inhibitory compounds

**DOI:** 10.1101/2021.01.18.427060

**Authors:** Thamiris Guerra Giacon, Gabriel Caetano de Gois e Cunha, Kevy Pontes Eliodório, Ricardo Pinheiro de Souza Oliveira, Thiago Olitta Basso

## Abstract

Second generation (2G) ethanol is produced through the use of lignocellulosic biomass. However, the pretreatment processes generates a variety of molecules (furan derivatives, phenolic compounds and organic acids) that act as inhibitors of microbial metabolism, and thus reduce the efficiency of the fermentation step in this process. In this context, the present study aimed to investigate the effect of furan derivatives on the physiology of lactic acid bacteria (LAB) strains that are potential contaminants of ethanol production. Homofermentative and heterofermentative strains of laboratory LAB and isolated from first generation ethanol fermentation were used. LAB strains were challenge to grow in the presence of furfural and hydroxymethyylfurfural (HMF). We found that the effect of HMF and furfural on the growth rate of LAB is dependent of the metabolic type, and growth kinetics in the presence of these compounds is enhanced for heterofermentative LAB, whereas is inhibitory to homofermentative LAB. Sugar consumption and product formation were also enhanced in the presence of furaldehydes in heterofermentative LAB, that displayed an effective detoxification kinetics when compared to the homofermentative LAB. This knowledge is important because LAB can be explored both within the scope of bio-detoxification, being applied before the fermentation.

**Key points:**

‐ Heterofermentative LAB presented the ability to decrease the concentrations of furfural and HMF
‐ LAB can be used in the bio-detoxification to remove the inhibitors before fermentations
‐ The presence of furan derivatives had a growth stimulus observed in heterofermentative LAB

## Introduction

The recent global economic development enhanced the demand for alternative energy resources worldwide, majorly due to the known drawbacks of fossil fuels, such as its high price, unsustainable and non-renewable feedstock, and global warming. On the other hand, one of the possible candidates to provide alternative energy is second-generation biofuels, which are produced from cheap and abundant plant biomass residues (Mood et al., 2013; Basso et al., 2013).

The second-generation (2G) biofuels are those produced through lignocellulosic materials, for instance, bioethanol, biodiesel, and biogas generated from these renewable materials. Among them, bioethanol is proving to be energetically and environmentally worthwhile when compared to the traditional first-generation ethanol. Furthermore, bioethanol from lignocellulosic materials suits the shortcoming of the first-generation by not applying edible feedstock sources (Aditiya et al., 2016).

The lignocellulosic biomass generally contains over 70% of carbohydrates in the form of cellulose and hemicelluloses in its composition, which may serve as a substrate for ethanol production (Klinke, et al., 2004). Cellulose is a polymer formed of glucose units whereas hemicellulose is a polymer formed of various units of xylose, arabinose, mannose, galactose, and glucose, which varies in composition depending on biomass (Bobleter, 1994; Fan et al. 1982). Associated with these carbohydrates is lignin in varying proportions, depending on the raw material source. Lignin is an amorphous polyphenolic compound with undefined molecular weight, predominantly composed of p-coumaryl alcohol, coniferyl alcohol, and synaphyl alcohol (Rubin, et al. 2008). Although the large content of carbohydrates, there are many chemical and physical barriers in lignocellulosic biomass that make it difficult for cellulose and hemicellulose to be available, requiring a pretreatment stage to make sugars easily fermentable by yeasts in ethanol production stage (Alvira, et al., 2010).

The main objectives of the pretreatment stage are to alter the lignin-hemicellulose-cellulose complex. The deconstruction of the complex reduces the crystallinity and increases its porosity and surface area, thus making it more accessible to the enzymatic hydrolysis reaction (Cardona, et al., 2010). These treatments can be physical, chemical, physicochemical, or biological. Most of these pretreatments, due to their severity, generate large amounts of inhibitory compounds to the metabolism of microorganisms applied in bioethanol production. The nature and concentration of these inhibitors are extremely affected by the adopted process and its operating conditions, such as temperature, time, pressure, pH, and presence of catalysts (Klinke, et al., 2004; van Maris et al., 2006). The inhibitors can be divided into 3 groups: furan derivatives, phenolic compounds, and organic acids. These compounds can severely affect the growth of microorganisms through DNA mutations, membrane disruption, intracellular pH decrease, among others (Chandel, et al., 2013).

Furans are formed mainly during pretreatments involving extremely acidic pH conditions due to the degradation of pentoses (2-furfural) and hexoses (5-hydroxymethylfurfural) (Dunlop, 1948). The concentration of furan aldehydes in lignocellulosic hydrolysates can range from 0.5 to 11 g.L^-1^ (Almeida et al., 2007). Aldehydes are chemically reactive and can form products with many classes of biological molecules. Several potential mechanisms for the toxicity of aldehydes have been explored, including damage from chemical reactivity, direct inhibition of glycolysis and fermentation, and plasma membrane damage (Zaldivar and Martinez, 1999). This class of inhibitors has been found to inactivate the cell replication that reduces the growth rate and the cell mass yield on ATP, volumetric growth rate, and specific productivities. At low concentrations, furans can have a positive effect on cell growth, although these compounds inhibit both cell growth and ethanol production rate by decreasing membrane permeability at higher concentrations (Taherzadeh, et al., 1999).

Adaptation of microorganisms on high furfural concentration has been found a successful option to overcome the negative effect of furfural on growth (Chandel, et al., 2011). Several enzymes have also been studied in yeast and bacterial cells, including NADH- and NADPH-dependent aldehyde reductases that can convert these compounds to the corresponding less inhibitory alcohols (Heer et al., 2009).

Since the formation of inhibitory by-products is not easily prevented economically at an industrial scale, it is often preferred to remove inhibitors before fermentation. Several types of detoxification strategies, such as physical (membrane-mediated detoxification, evaporation), chemical (overliming, calcium hydroxide, neutralization, ion exchange resins, activated charcoal column, and extraction with ethyl acetate), biological (microbial and enzymatic), and in situ microbial detoxification, have been applied to remediate fermentation inhibitors. Biological methods of detoxification are in demand due to their simplicity, milder work conditions that avoid further use of chemicals, lower frequency of side reactions, and lower energy requirements (Parawira and Tekere, 2011). The main drawbacks of biological methods include slow reaction time, inhibitor specificity, and loss of fermentable sugars, which makes them unattractive to biorefineries (Tian et al., 2009).

Among the biological options, *Lactobacillus* spp. arises as one possible candidate as a result of its ability to transform furfural and HMF in less toxic compounds of furfuryl alcohol and 2,5-bis(hydroxymethyl)furan, respectively. This process is also known as “in situ-detoxification” (Boopathy et al., 1993). Lactic bacteria (LAB) can be classified as homo and heterofermentative.

Homofermentative LAB use the Embden-Meyerhof Parnas pathway for glucose catabolism and the pyruvate formed is reduced to lactic acid. Furfural and HMF are able to inhibit important metabolic enzymes, including alcohol dehydrogenase, aldehyde dehydrogenase, and pyruvate dehydrogenase.

Heterofermentative LAB converts glucose via the phosphoketolase pathway, resulting in an equimolar mixture of lactic acid and ethanol/acetic acid (Kandler, 1983). It has the ability to reduce furfural with NADH and NADPH using the furan derivatives as alternative electrons acceptors, enhancing regeneration of NAD+. This can result in increased bacteria growth while blocking the production pathways of lactic acid and increasing the acetate/ethanol pathway because furfural and HMF are acting as NAD and NADP recyclers and the acetate/ethanol production route becomes energetically more advantageous.

In this context, the aim of this research was to study the effect of furaldehydes from lignocellulosic hydrolysates on the physiology of different strains of lactic acid bacteria, divided into homo and heterofermentative metabolism and laboratory and industrial. strains. This knowledge is important because lactic acid bacteria can be explored both within the scope of bio-detoxification, being applied before the fermentation, as well as in the knowledge of the mechanisms used by these bacteria in the detoxification to be applied in other microorganisms, exploring genetic engineering.

## Material and methods

### Strains

The microorganisms used were 12 lactobacilli strains in initial screening experiments, five homofermentative (*Lactobacillus plantarum* CECT 221, *Lactobacillus delbrueckii, Lactobacillus plantarum* ESALQ 4, *Lactobacillus paracasei* LAB 4, *Lactobacillus paracasei* LAB 5) and seven heterofermentative (*Lactobacillus fermentum* DSM 20391, *Lactobacillus reuteri* ATCC 23272, *Lactobacillus fermentum* ESALQ 3, *Lactobacillus fermentum* ESALQ 5, *Lactobacillus fermentum* 1L-6-MRS, *Lactobacillus fermentum* 3L-2-M17, *Lactobacillus paracasei* LAB 2). Strain codes and sources are provided in Table 1.

**Table 1.** LAB strains

| Code | Strain | Isolation |
| --- | --- | --- |
| A | *Lactobacillus plantarum* CECT 221 | Food industry |
| B | *Lactobacillus delbrueckii* | Beer industry |
| C | *Lactobacillus plantarum* ESALQ 4 | Sugar cane fermentation |
| D | *Lactobacillus paracasei* LAB 4 | Sugar cane fermentation |
| E | *Lactobacillus paracasei* LAB 5 | Sugar cane fermentation |
| F | *Lactobacillus fermentum* DSM 20391 | Human oral cavity |
| G | *Lactobacillus reuteri* ATCC 23272 | Rat intestine |
| H | *Lactobacillus fermentum* ESALQ 3 | Sugar cane fermentation |
| I | *Lactobacillus fermentum* ESALQ 5 | Sugar cane fermentation |
| J | *Lactobacillus fermentum* 1L-6-MRS | Sugar cane fermentation |
| K | *Lactobacillus fermentum* 3L-2-M17 | Sugar cane fermentation |
| L | *Lactobacillus paracasei* LAB 2 | Sugar cane fermentation |

### Propagation and storage of microbial strains

Bacterial strains were grown in De Man, Rogosa & Sharpe medium (MRS) with glucose (20 g.L^-1^), peptone (10 g.L^-1^), meat extract (10 g.L^-1^), yeast extract (5 g.L^-1^), K_2_HPO_4_ (2 g.L^-1^), sodium acetate (5 g.L^-1^), tri-ammonium citrate (2 g.L^-1^), MgSO_4_.7H_2_O (200 mg.L^-1^), MnSO_4_.4H_2_O (50 mg.L^-1^), Tween 80 (1 mL.L^-1^), The pH was adjusted to 6 and temperature of cultivation to 37 °C. From the final volume obtained, 20% relative to it of glycerol was added and 2 mL aliquots were stored in a freezer at −80 °C.

### Inoculum preparation

The inoculum was prepared in a 50 mL conical tube containing 25 mL of MRS medium, where 200 µL of stock culture was added. The inoculum was grown for 24 h at 37°C.

### Semi-defined media supplemented with furanic derivatives

To better evidence the stress response with the presence of furanic compounds, the experiments were performed in semi-defined medium containing glucose (20 g.L^-1^), yeast extract (5 g.L^-1^), peptone (5 g.L^-1^), K_2_HPO_4_ (2 g.L^-1^), MgSO_4_ (0.2 g.L^-1^), and MnSO_4_ (0.01 g.L^-1^) (Basso et al.,2014). The pH was adjusted to 6 and temperature conditions were controlled at 37°C for 24 h.

### Screening of bacterial growth with furanic derivatives

Each of the strains were incubated into 96-well plate growth containing two concentrations of each of the furanic compounds studied. Furfural concentrations were 1.5 g.L^-1^(15.6 mM) and 2.5 g.L^-1^(26 mM) and HMF concentrations were 2 g.L^-1^ (15.85mM) and 4 g.L^-1^(31.7mM). Control cultures were performed at the same time without the supplementation of inhibitory compounds. Triplicate cultures were carried out for each treatment. Concentrations were based on previously reported studies on literature data (Cola et al., 2020; van der Pol et al. 2014).

The growth was performed in semi-defined medium were the OD_600_ was evaluated every 15 minutes with mean value of five reads per well in the microplate reader Tecan Infinite M200, from which the growth curve was calculated to estimate the specific maximum velocity (µ).

### Kinetic assays

The kinetic assays were performed in 50 mL conical tubes with 30 mL of semi-defined medium with the highest concentration of each inhibitor (HMF 4g .L^-1^ and furfural 2.5g.L^-1^). Control cultures were not supplemented with either inhibitor. Triplicate cultures were carried out for each treatment. The experiments were carried out at 37°C until the sugar level in the medium was finished. Aliquots were taken from time to time to measure OD_600_ and to track the sugar consumption and the metabolite production.

### Maximum specific growth rate and lag phase estimation

The maximum specific growth rate (*µ*_*max*_) and the lag phase (*λ*) were determined by carefully applying the empirical sigmoidal model of Morgan-Mercer-Flodin (Eq.1) (Tjørve, 2003) to the natural logarithm of the bacterial count (y=ln(N)), determined as OD_600_ reads.

The corresponding equations to calculate the microbiological parameters (*µ*_*max*_ *and λ*) based on model parameters (A, b, and n) are given in Table 2. The complete mathematical approach and theoretical background are described in detail by Longhi et al. (2017).

**Table 2.** Equations to *µ*_*max*_ *and λ based on* parameters (A, b, and n) of Morgan-Mercer-Flodin model

| Morgan-Mercer-Flodin model |  |
| --- | --- |
| $y(t) = y_0 + \frac{A \cdot t^n}{b + t^n}$ | (Eq.1) |
| $\mu_{max} = \frac{A \cdot (n-1)^{\left(\frac{n-1}{n}\right)} \cdot (n+1)^{\left(\frac{n+1}{n}\right)}}{4 \cdot n \cdot \sqrt[n]{b}}$ | (Eq. 2) |
| $\lambda = \sqrt[n]{b} \cdot \left(\frac{n-1}{n+1}\right)^{\left(\frac{n+1}{n}\right)}$ | (Eq. 3) |

The parameters of the sigmoidal model are the upper asymptote parameter (A) and the shape parameters (b and n).

The fitting of the mathematical model to the experimental data was assessed in the optimization toolbox of MATLAB R2015b software (MathWorks, Natick, USA). The *lsqcurvefit* function was applied using a non-linear least-squares method and the trust-region reflective Newton algorithm with as initial value of parameters selected by experimental data observation. The Adjusted Coefficient of Determination (*R* ^*2*^ _*adj*_) and the square sum of the residual were used to evaluate the quality of the fitting procedure to the experimental data.

### Analytical methods

Metabolite samples were immediately centrifuged and stored at −20 °C until analysis. Sugar consumption and the metabolite production were analyzed using HPLC Prominence (Shimadzu) with ion exclusion column Aminex ^®^HPX-87H (300×7.8mm×9 μm) (Bio-RAD), isocratically eluted at 60 °C with 5 mM sulfuric acid flow rate of 0.6 mL.min-1. The total run time was 50 min, refractive index detector was used.

## Results

### The effect of HMF and furfural on the growth rate of lactic acid bacteria is dependent of the metabolic type

The effect of HMF and furfural was studied in lactic acid bacteria (LAB) displaying homo- and heterofermentative metabolism. According to the results obtained in the microplate reader cultivations, it was observed that the inhibitory compounds affected differently the two groups of LAB. The two inhibitory furan-derivative compounds had a positive effect on the growth of the heterofermentative LAB (Fig. 1), increasing its maximum specific growth rate by up to 2.4 times in comparison to the control condition (cultivation media in the absence of the inhibitory compounds). In addition, a positive effect for both compounds in the elongation of the lag phase was also observed in the heterofermentative LAB, in which a decrease in the time required to reach the exponential phase was noticed as compared to the control condition (Fig. 2). On the other hand, in the homofermentative LAB, the complete opposite effect was observed, since their growth rates were inhibited by the furan-derivatives as compared to the control, as shown in Fig. 1. Likewise, homofermentative LAB also had a negative effect on the lag phase duration, when cells were cultivated in the presence of the two inhibitors (Fig. 2).

**Figur 1.**
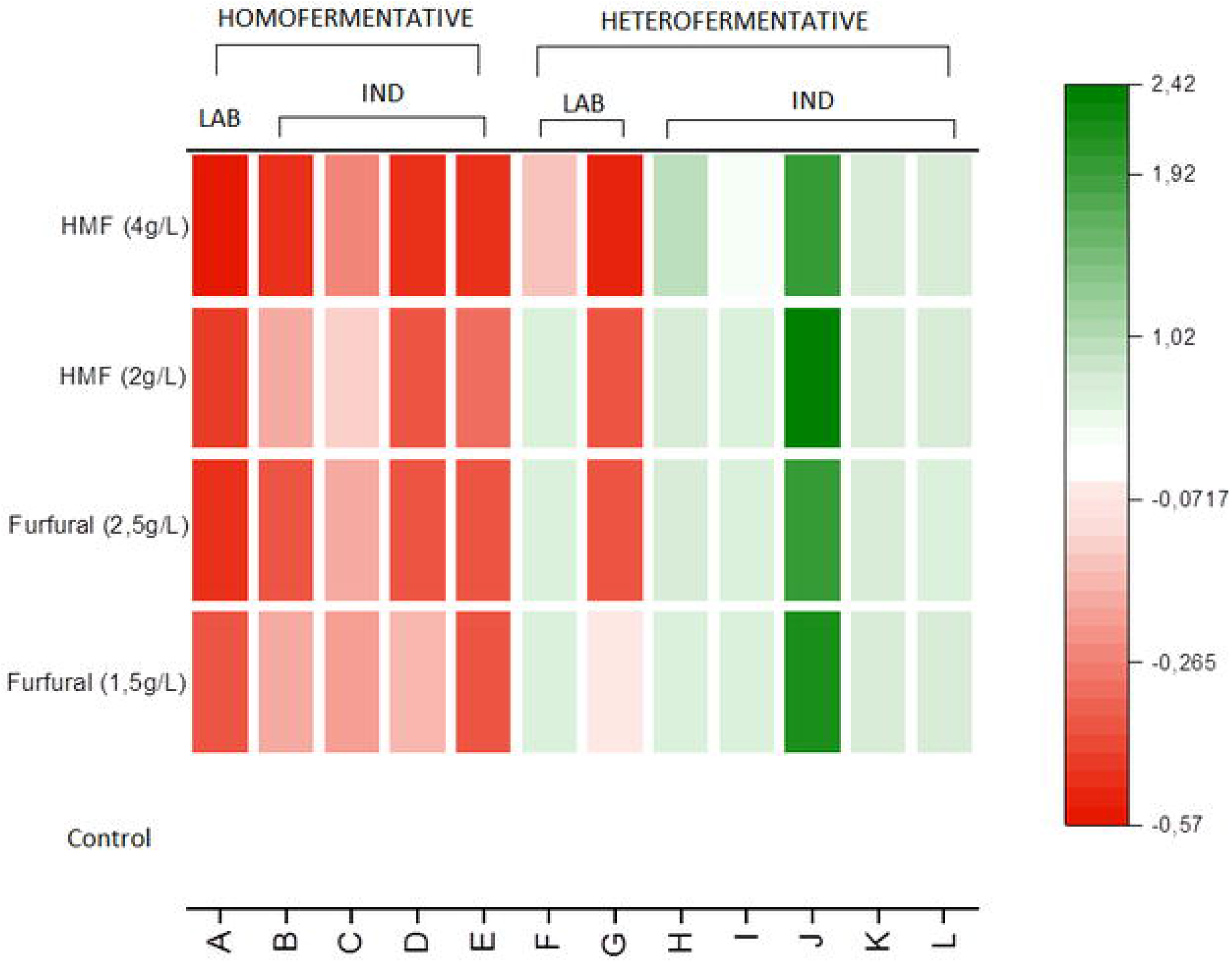

**Figur 2.**
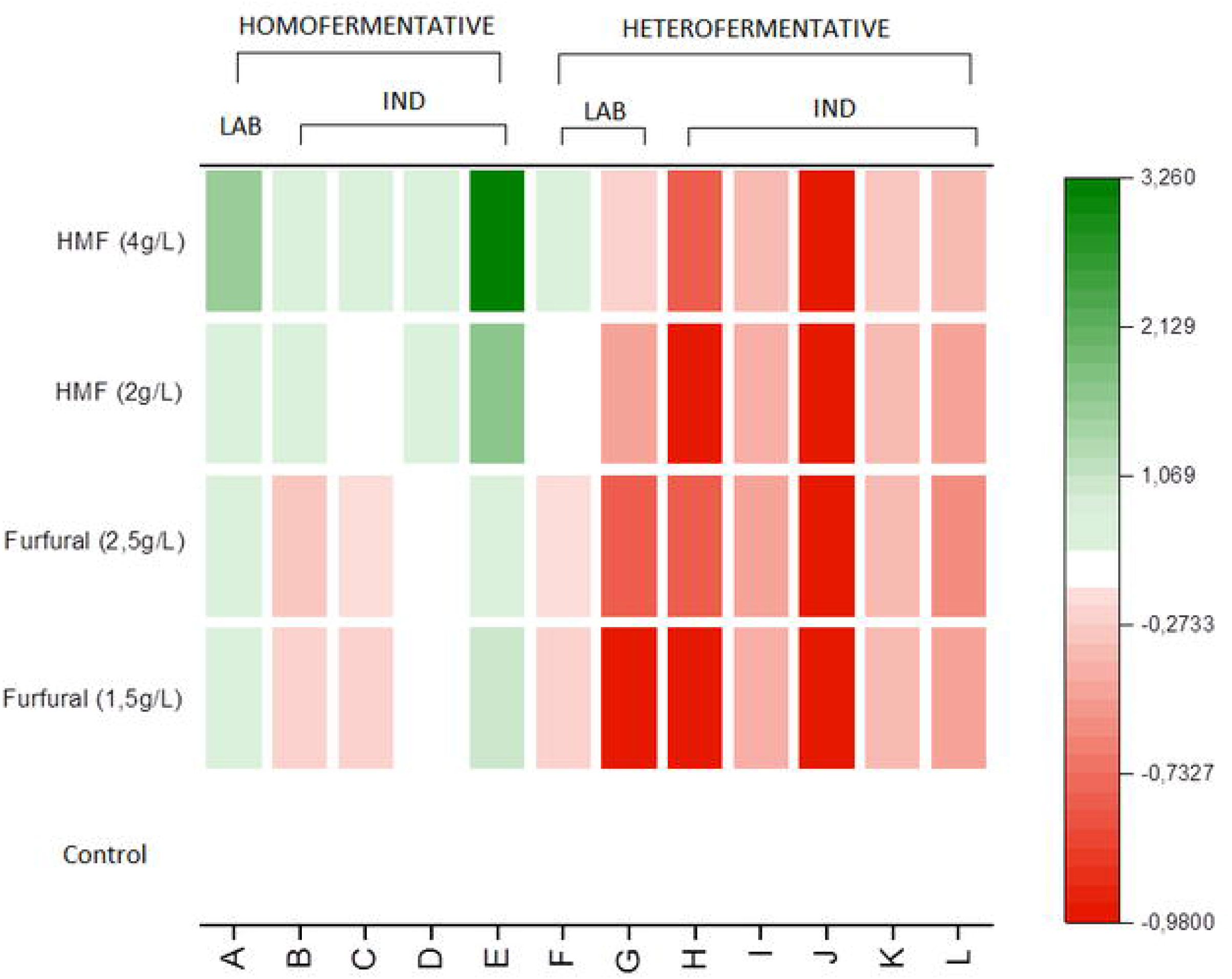

When comparing laboratory and industrial strains, it was observed that the growth performance (evaluated by the maximum specific growth rate and the elongation of the lag phase) of the laboratory heterofermentative LAB strains were less stimulated or partially inhibited than in the industrial LAB strains. Comparing laboratory and industrial homofermentative LAB strains, it was not possible to observe a pattern for these two parameters that could be used to differentiate them. The only exception was the fact that in all conditions tested the decrease in the growth rate of the homofermentative laboratory strain was more pronounced than the industrial strains.

### Growth kinetics in the presence of HMF and furfural is only enhanced for heterofermentative lactic acid bacteria

As a follow-up, the growth kinetics was further investigated in two representative LAB strains using shake-flask cultures with the same semi-defined media supplemented with the two furan derivatives separately at the highest concentration tested. For that purpose, a representative homofermentative (*Lactobacillus plantarum* ESALQ 4) strain, and a representative heterofermentative (*Lactobacillus fermentum* ESALQ 3) strain were investigated in these conditions. The measurement of growth give us reliable and sensitive information for the characterization of toxic compounds and conditions that adversely affect microbial cells (Franden et al., 2009)

As observed in the general growth screening described above in microplate cultures, the heterofermentative LAB showed faster growth kinetics and shorter lag phase in the presence of both inhibitors as compared to their absence (Fig. 3a). The exponential phase starts almost with 5 h of cultivation in the control kinetics and could be evidenced in 2 h in furfural presence. This effect is less pronounced in the presence of HMF.

**Figur 3.**
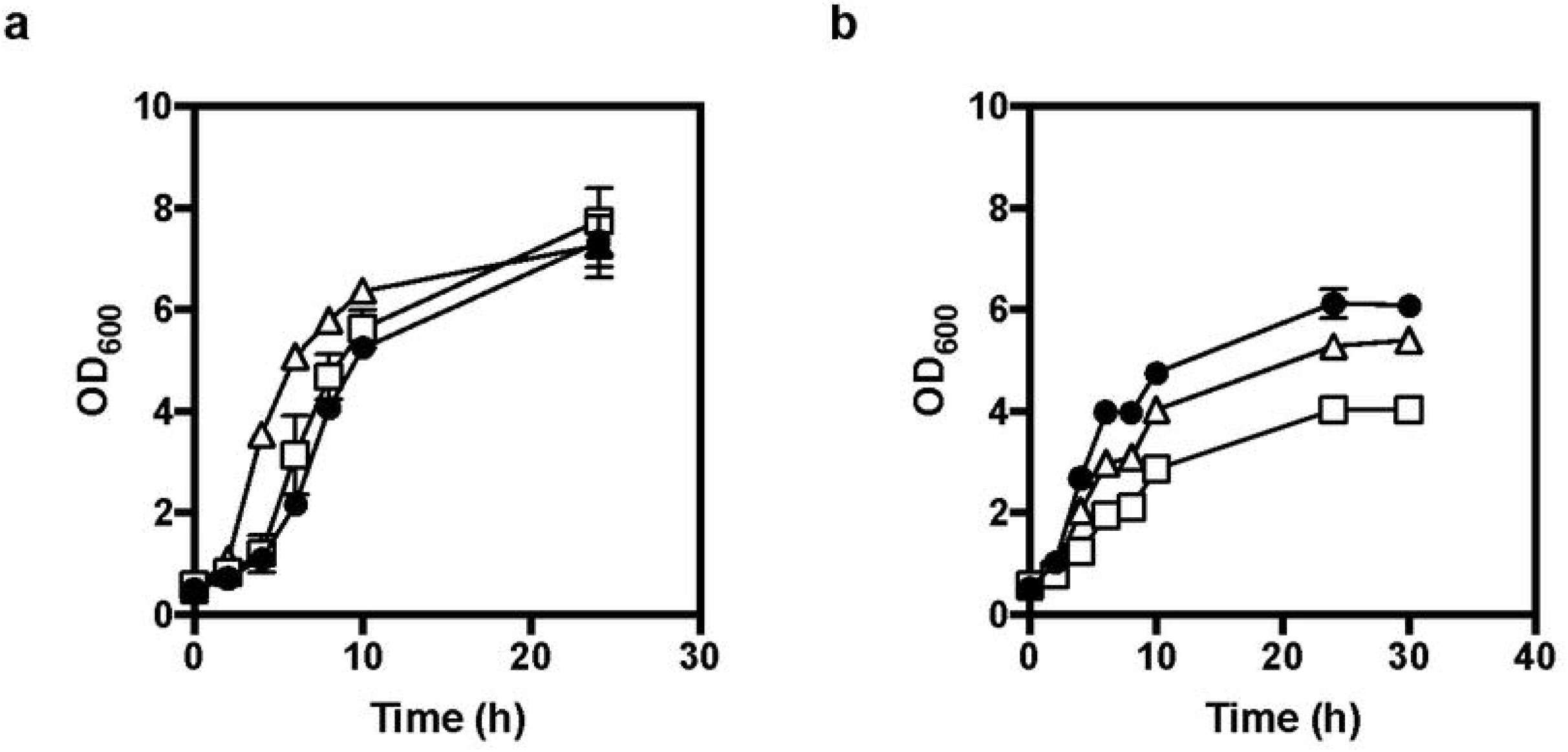

On the other hand, the growth kinetics in the presence of both inhibitors was inhibited as compared to the control condition (absence of furan derivatives) in homofermentative LAB. Apparently, in the concentrations tested, HMF seemed to be more detrimental to the homofermentative strain than furfural (Fig. 3b).

### Sugar consumption and product formation is enhanced in the presence of furaldehydes in heterofermentative lactic acid bacteria

Heterofermentative LAB normally present slow growth kinetics on glucose that is caused by the low activity of the ethanol pathway in the reoxidation of the extra two NADH (Maicas et al.,2002). In the presence of the furan inhibitors, we observed an enhanced sugar consumption rate, and we noticed a deviation towards the formation of acetate and ethanol with a concomitant decrease in lactate production when compared to the control condition (Fig. 4, Table 3). It seems this observation is caused by the fact that furfural and HMF are promoting the reoxidization NAD+ and NADP+, respectively. In addition, as they do not need to use the ethanol route to reoxidize de NADH, the acetyl-P can be used by ATP synthesis and the acetate production route becomes energetically more advantageous (Ganzle, 2015).

**Table 3.** Conversion yields (in g.g^-1^) homofermentative LAB strain cultivated in media containing furfural and HMF. Values are expressed as means (n = 3). Different letters in the same column indicate significant difference (P ≤ 0.05).

| Condition | Y<sub>sx</sub> | Y<sub>sl</sub> |
| --- | --- | --- |
| Control | 0.100±0.004 (A) | 1.148±0.039 (A) |
| HMF (4.0 g L<sup>-1</sup>) | 0.089±0.005 (B) | 1.156±0.038 (A) |
| Furfural (2.5 g L<sup>-1</sup>) | 0.093±0.002 (AB) | 1.132±0.009 (A) |

**Table 4.** Conversion yields (in g.g^-1^) heterofermentative LAB strain cultivated in media containing furfural and HMF. Values are expressed as means (n = 3). Different letters in the same column indicate significant difference (P ≤ 0.05).

| Condition | Y<sub>sx</sub> | Y<sub>sl</sub> | Y<sub>se</sub> | Y<sub>sa</sub> |
| --- | --- | --- | --- | --- |
| Control | 0.093±0.007 (A) | 0.854±0.013 (A) | 0.113±0.004 (A) | 0.032±0.0007 (A) |
| HMF (4.0 g L<sup>-1</sup>) | 0.087±0.007 (AB) | 0.592±0.016 (B) | 0.175±0.015 (B) | 0.101±0.008 (B) |
| Furfural (2.5 g L<sup>-1</sup>) | 0.076±0.004 (B) | 0.579±0.007 (B) | 0.221±0.004 (C) | 0.068±0.0004 (C) |

**Figur 4.**
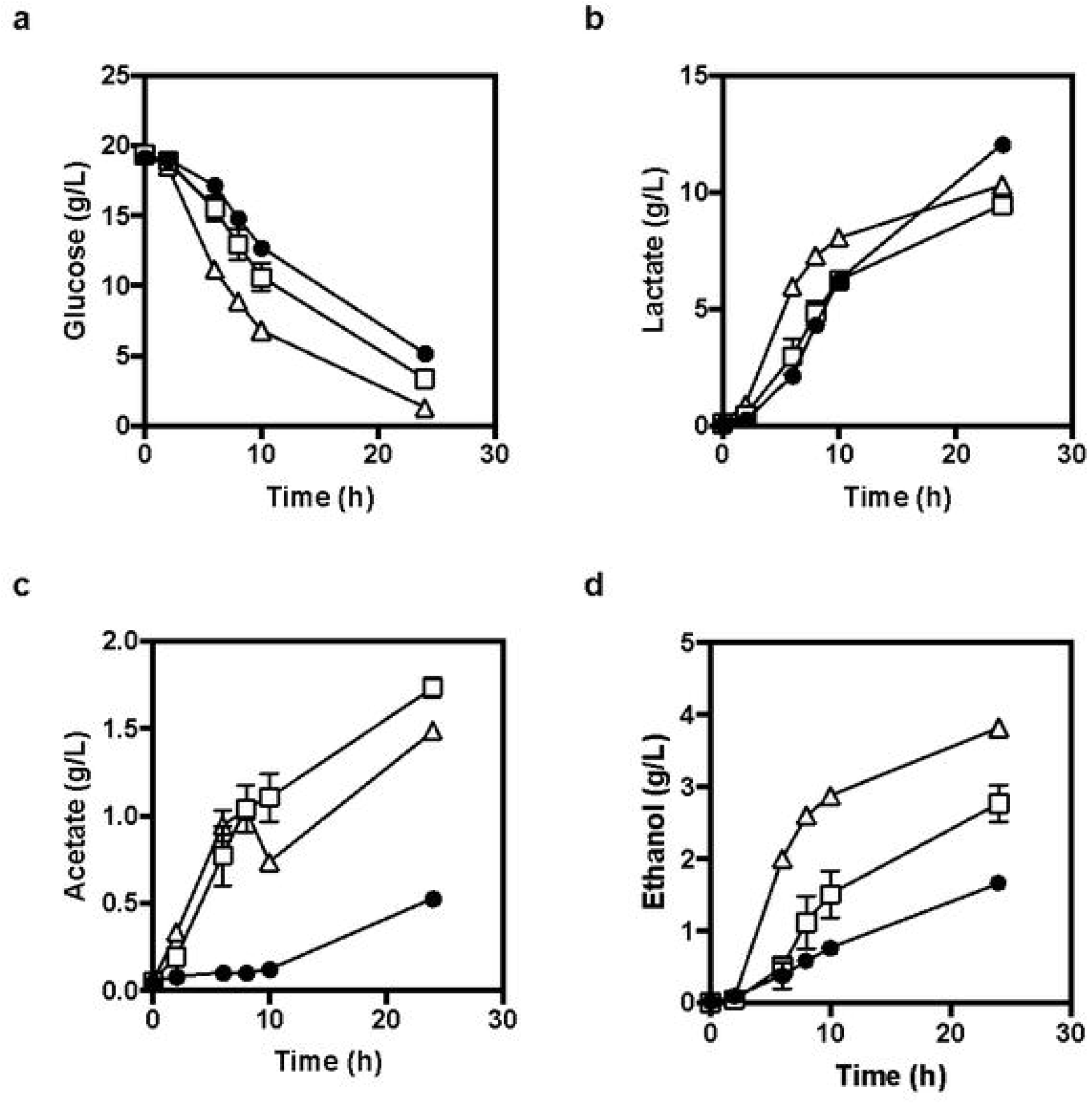

In cultures with the homofermentative LAB, the opposite behavior was observed; sugar consumption rate decreased in the presence of inhibitors and it was also possible to notice a decrease in lactate production kinetics (Fig. 5). This is probably because this strain was not able to detoxify the media, and consequently, furfural and HMF posed a deleterious effect on the cells (Fig. 6).

**Figur 5.**
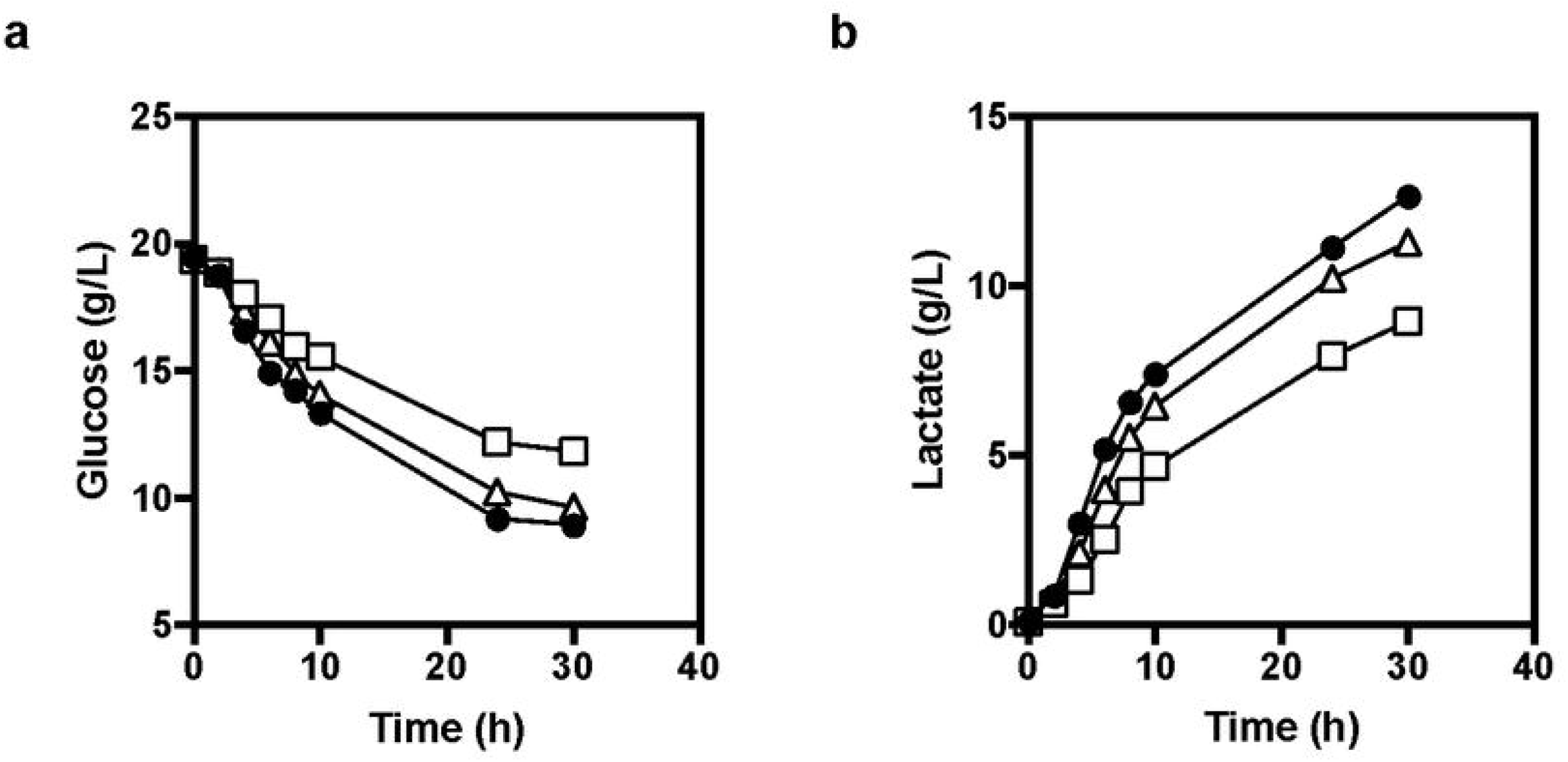

**Figur 6.**
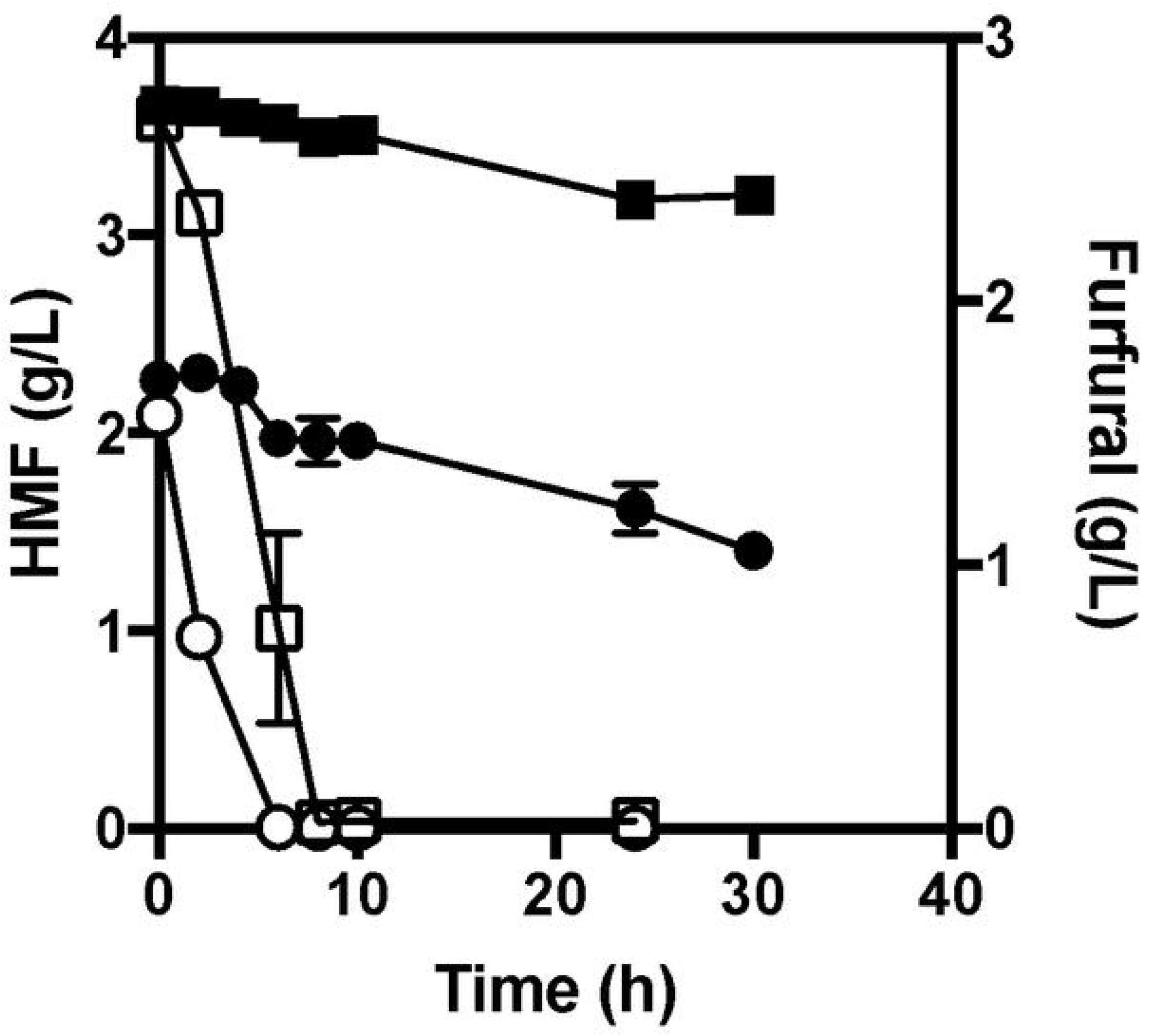

Furfural and HMF acted as an enzyme inhibitor like have been shown in previous works, since they can inhibit important metabolic enzymes. In *Saccharomyces cerevisiae*, including alcohol dehydrogenase, aldehyde dehydrogenase, and pyruvate dehydrogenase, thus decreasing viability and growth and increasing lag phase duration (Modig et al., 2002). Bai (2015) reported the inhibition in *Sporolactobacillus inulinus* of four key enzymes of the lactic acid pathway by the furanic derivatives, with a decrease of almost 80% in the lactate dehydrogenase activity in the presence of 5g.L^-1^ of furfural. HMF had a weaker inhibitory effect than furfural, which can explain the decrease in lactic acid production in both LAB tested.

### Detoxification of furaldehydes is very effective in heterofermentative lactic acid bacteria

Homofermentative bacteria dissimilate hexoses through glycolysis, where fermentation of 1 mol of hexose results in the formation of 2 mol of lactic acid and 2 mol of ATP. In comparison, heterofermentative bacteria present another active pathway (Kandler and Weiss, 1986) and hexoses are converted to equimolar amounts of lactic acid, ethanol or acetate, and carbon dioxide, yielding 1 mol of ATP per mol of hexose fermented (Cogan and Jordan, 1994). With the conversion of acetyl phosphate to acetate instead of ethanol, an additional ATP can be produced. Then, the regeneration of surplus NAD+ must be achieved through an alternative electron acceptor.

In the heterofermentative LAB, it was possible to see the complete depletion of HMF and furfural which we hypothesized that were converted to furfuryl alcohol and furandimethanol (FDM), respectively (Fig. 6). Previous studies indicate that yeast and bacteria strains were able to reduce furfural and HMF to their corresponding alcohols, as reported for *Lactobacillus reuteri* (van Niel, 2012), *Saccharomyces cerevisiae (*Liu et al., 2011), *and E. coli* (Jozefczuk, 2010) in which. These degradation products are of lower toxicity to microorganisms compared to their aldehyde precursors (Liu et al., 2011).

Heterofermentative *L. fermentum* seems to convert HMF at a slower rate as compared to furfural, which may be attributed to a lower cell membrane permeability of HMF when compared to furfural (Larsson et al., 1999). Therefore, under the oxygen-limited conditions that the experiments were performed, furan derivatives might have been reduced to their corresponding alcohols. In this way, such inhibitors seemed to be important co-substrates for heterofermentative lactobacilli, as opposed to homofermentative strains (Fig. 8).

## Discussion

According to the results it was observed that the two inhibitory furan-derivative compounds had a positive effect on growth of the heterofermentative LAB. Even when added in high concentrations to a culture in an early exponential growth phase, the compounds seem to enhance growth performance in the heterofermentative LAB. In this case, we hypothesized that the heterofermentative bacteria has the ability to reduce furfural with NADH and NADPH using furfural and HMF as alternative electrons acceptors. The reduction of furfural is preferentially dependent on NADH, and the reduction of HMF has been mainly associated with the consumption of NADPH (Wahlbom and Hahn-Hägerdal, 2002). This behavior was also observed in *Lactobacillus reuteri* by van Niel (2012) and other several microorganisms like *Saccharomyces cerevisiae* (Liu et al., 2011), *Escherichia coli* (Gutierrez et al., 2002) and other enteric bacteria (Boopathy et al., 1993). Also, it was observed the decreased in lag phase in heterofermentative LAB, this was also observed in *Saccharomyces cerevisiae* by Liu (2009) because of the alcohol and aldehyde dehydrogenases upregulation that caused this decrease in the lag phase elongation and an increase in furan tolerance.

It is possible that the presence of HMF or furfural in the medium may have enhanced glycolysis via regeneration of NAD^+^, because NADH may be involved in the reduction of these furans to their corresponding alcohols (furfuryl alcohol and HMF alcohol). Therefore, in view of the enhanced regeneration of NAD^+^ in the presence of furans, glycolytic flux might have been enhanced in the cultures of the heterofermentative *L. fermentum* strain. The alcohol form reduced from the aldehyde form appeared to not affect bacteria fermentation and the accumulated detoxification products in the medium did not affect the final lactate production.

In homofermentative LAB many metabolic processes may be significantly altered and delayed in the presence of these inhibitors (Vertes et al., 2011). Furfural and HMF may inhibit glycolysis pathway and hexokinases responsible for phosphorylation of six-carbon sugars. Furfural is reported to cause cell membrane damage and to inhibit the activity of various glycolytic enzymes, such as hexokinase and glyceraldehyde-3-phosphate dehydrogenase (Almeida et al., 2007). Therefore, pyruvate production by this pathway would be depleted, leading consequently to a decrease in the lactic acid concentration in the homofermentative LAB. The deleterious effects observed on the growth kinetics in the homofermentative strains are sought to be due to enzyme inhibition and to damage in the cell membrane, that are exacerbated by the fact that these compounds are not metabolized by the homofermentative strain (Taherzadeh and Karimi, 2011).

Nevertheless, it is unclear to which extent furanic compounds are truly metabolized. In several reports, it is merely established that the furanic aldehydes have disappeared, without mention of the metabolic pathways of the corresponding alcohols or carboxylic acids. Therefore, all the different forms of the furanic compound (alcohol, aldehyde, and carboxylic acid) should be carefully monitored in order to establish whether the furanic aldehydes are actually metabolized or only transformed into a less toxic form (Wierckx et al. 2010; Wierckx et al., 2011).

## Conclusions

The heterofermentative bacterium presented the ability to decrease the concentrations of furfural and HMF in the fermentation medium, with simultaneous lactic acid production. Low concentrations of these compounds present in the sugarcane bagasse hemicellulosic liquor did not have inhibitory effects on the lactic acid production. The approach of bio-detoxification of the fermentation broth dispenses a hemicellulosic liquor detoxification process prior to fermentation and expands the possibility for lactic acid production from second-generation feedstock.

A better understanding of the genetic mechanisms and biochemical pathways responsible for inhibitor response in bacteria may allow the development of genetically engineered novel strains to withstand major inhibitors generated from biomass pretreatment.

## Acknowledgements

This work was financially supported in part by CNPq and FAPESP (2019/13826-0)

## Availability of data and material

Not applicable

## Code availability

Not applicable

## Ethical approval

This article does not contain any studies with human participants or animals performed by any of the authors

